# Acute anaphylactic and multiorgan inflammatory effects of Comirnaty in pigs: evidence of spike protein mRNA transfection and paralleling inflammatory cytokine upregulation

**DOI:** 10.1101/2025.06.07.658379

**Authors:** László Dézsi, Gábor Kökény, Gábor Szénási, Csaba Révész, Tamás Mészáros, Bálint A. Barta, Réka Facsko, Anna Szilasi, Tamás Bakos, Gergely T. Kozma, Attila B. Dobos, Béla Merkely, Tamás Radovits, János Szebeni

## Abstract

**Background and Purpose:** Rare but serious adverse events (AEs) associated with mRNA-lipid nanoparticle (LNP) COVID-19 vaccines, such as Comirnaty and Spikevax, include anaphylaxis and acute multiorgan inflammatory syndrome. The mechanisms of these acute innate immune responses remain poorly understood. This study aimed to investigate these effects using an amplified porcine model.

**Experimental Approach:** Naïve and anti-PEG antibody-sensitized pigs were intravenously injected with Comirnaty. Acute anaphylactic responses were assessed through hemodynamic monitoring, hematological changes, and plasma inflammatory markers. Multiorgan inflammatory effects were evaluated by RT-qPCR detection of spike protein (SP) mRNA uptake and inflammatory cytokine gene expression in seven organs over 6 hours. Histopathology and immunohistochemistry were also performed.

**Key Results:** Severe cardiopulmonary distress developed within minutes of repeated intravenous injections of Comirnaty. RT-qPCR revealed predominant SP mRNA accumulation in liver and PBMCs, with spleen, kidney, lymph nodes, heart, and brain also affected in 40–90% of animals. Repetitive PBMC transfection showed reversible mRNA peaks at 15 minutes, followed by rapid decay. Expression of proinflammatory cytokines (IL-1RA, CXCL10, TNF-α, CCL2) paralleled mRNA uptake, suggesting a causal relationship. Cytokine profiles varied by organ (e.g. IL-1RA in kidney, CCL2 in PBMCs).

Histologic abnormalities and SP immunopositivity were noted in kidney, heart, and brain. Booster injections replicated PBMC transfection kinetics observed at first dose.

**Conclusions and Implications:** This porcine model reveals systemic anaphylactic reactivity and multiorgan SP mRNA transfection following intravenous injection of mRNA-LNPs, resulting in organ-specific inflammatory responses. These findings provide mechanistic insights into two rare vaccine-related acute AEs and may support future efforts to improve the safety of the mRNA-LNP platform technology.

## 1. INTRODUCTION

During the COVID-19 mRNA vaccination campaign, the rare but globally recognized broad spectrum of adverse events (AEs) linked to Pfizer’s Comirnaty and Moderna’s Spikevax drew intense scientific and public attention. Emerging evidence and recent policy changes have acknowledged the clinical significance of these AEs, prompting a shift toward risk-based vaccination recommendations and renewed efforts to elucidate the underlying mechanisms of the phenomenon. These AEs can be acute or chronic. Among the acute reactions, anaphylaxis (Barta et al., 2024; Dezsi et al., 2022; Kozma et al., 2023; Paul et al., 2022; Shimabukuro et al., 2021; Song et al., 2025) and multisystem inflammatory syndrome (Salzman et al., 2021; Yousaf et al., 2023) are among the most serious. These conditions share several clinical features, such as the rapid-onset systemic inflammation, multiorgan involvement, cardiovascular instability, and potential fatality in severe cases. Nevertheless, their immunopathogenesis substantially differ, most notably, anaphylaxis represents an overactivation of the humoral arm of the innate immune system, primarily manifested by anaphylatoxin release, whereas multisystem inflammatory syndrome is driven by the cellular arm, with predominant cytokine release.

Although the overall incidence of mRNA-vaccine-induced anaphylaxis is low (up to 0.04%) (Anis et al., 2022; Blumenthal et al., 2021; Shimabukuro et al., 2021; Szebeni, 2025; Warren et al., 2021), it was estimated to be >60-fold higher compared to that of flu vaccines (Szebeni, 2025a). Multisystem inflammatory syndrome in children and adults are even rarer, occurring in a few per million (CDC, 2025; Hamad Saied et al., 2023; Joob & Wiwanitkit, 2023; Salzman et al., 2021; Vogel et al., 2021; Yousaf et al., 2022; Yousaf et al., 2023), and, unlike anaphylaxis, epidemiological analysis of prevalence or incidence have not been reported after any other vaccines, only a few case reports link multisystem inflammatory syndrome to adenovirus-based COVID-19 vaccines (Bova et al., 2022). Multisystem inflammatory syndrome is characterized by inflammation across multiple organ systems, including the heart. The underlying mechanism is hypothesized to involve innate immune activation; however, details of this process is poorly understood. Among other unanswered questions, it is not known whether the inflammation is caused by the SP, or the LNP, or both in an additive or synergistic manner. The pig model applied here enables in-depth analysis of these questions, and hence advance the understanding, prediction and prevention of these rare but severe vaccine complications.

## 2. METHODS

### 2.1. Pig studies

Landrace pigs were obtained from the Research Institute for Animal Breeding, Nutrition and Meat Science of the Hungarian University of Agriculture and Life Sciences (Herceghalom, Hungary) and SPF Danbred pigs from Sano-Pikk Ltd., (Zsámbék, Hungary). The study involved 11 (+2 SPF) female and castrated male pigs in the 22–32 kg size range. They underwent a 1-week acclimation period in the experimental large animal housing facility before starting the experiments. Pigs were given standard laboratory pig diet and water *ad libitum*. All procedures were performed in accordance with ARRIVE guidelines and the guidelines of the Council Directive of the European Communities (86/609/EEC) and approved by the Pest County Government Office, under permission number: PE/EA/843-7/2020. SPF female and castrated male Danbred pigs (X-Ykg BW) were obtained from Sano-Pikk Ltd., Zsámbék, Hungary, underwent a 1-week acclimation period in the experimental large animal housing facility before starting experiments. Pigs were given standard laboratory pig diet and water *ad libitum*.

### 2.2. Immunization against liposomal PEG

In the first stage animals were immunized against liposomal PEG by injecting them with liposomal doxorubicin (Doxebo). The preparation and characteristics of Doxebo were described earlier in detail (Kozma et al., 2019). In brief, the freeze-dried lipid components of Doxil were hydrated in 10 mL sterile pyrogen-free normal saline by vortexing for 2−3 min at 70°C to form multilamellar vesicles (MLVs). The MLVs were downsized through 0.4 and 0.1 μm polycarbonate filters in two steps, 10 times through each, using a 10 mL extruder barrel from Northern Lipids (Vancouver, British Columbia, Canada) at 62 °C. Liposomes were suspended in 0.15 M NaCl/10 mM histidine buffer (pH 6.5). The size distribution (Z-average: 81.17 nm) and phospholipid concentration of Doxebo (12.6 mg/mL) were determined as described earlier (Kozma et al., 2019). After taking preimmune blood samples ani-PEG immunization was done by infusing 0.1 mg PL/kg Doxebo via the ear vein (suspended in 20 mL of saline) at a speed of 1 mL/min. The animals were then placed back into their cages until the 2nd blood sampling 7-10 days later, to screen for anti-PEG Ab induction. From that time the animals showing seroconversion were subjected to the “CARPA induction” protocol with Comirnaty. One control animal was sham-immunized with PBS but was handled identically to the Doxebo-immunized pigs.

### 2.3. Treatment of pigs with Comirnaty

We report in this study data from 3 types of pig experiments. In one, pigs were treated with three consecutive intravenous injections of Comirnaty, each containing 0.3 ug mRNA/kg (~3X human dose). Thereafter, the animals were subjected to the “standard” CARPA protocol (Szebeni, 2024) as follows. After sedation in the porcine confinement area with an i.m. injection of 25 mg/kg ketamine (Gedeon Richter Plc. Budapest, Hungary) and 0.3 mg/kg midazolam (Kalceks AS, Riga, Latvia), the animals were carefully transported into the laboratory. Anesthesia was induced with a propofol bolus through an auricular vein. Airways were secured by inserting an endotracheal tube. Unless otherwise specified, animals were allowed to breath spontaneously during the experiments. Controlled ventilation was applied when continuous measurement of hemodynamic parameters necessitated invasive surgical interventions. Surgery was done after povidone iodine (10%) disinfection of the skin. To measure the pulmonary arterial pressure (PAP), a Swan-Ganz catheter (Arrow AI-07124, 5 Fr. 110 cm, Teleflex, Morrisville, NC, USA) was introduced into the pulmonary artery via the right internal jugular vein. A Millar catheter (SPC-561, 6 Fr. Millar Instruments, Houston, TX, USA) was placed into the left femoral artery to record the systemic arterial pressure (SAP). Additional catheters were introduced into the left external jugular vein for drug administration, into the left femoral vein for venous blood sampling, and into the right common carotid artery for arterial blood gas analysis. The latter was executed with a Roche COBAS B221 benchtop analyzer (Roche Diagnostics, Rotkreuz ZG, Switzerland). Hemodynamic data were collected using instruments from Pulsion Medical Systems, and Powerlab, AD-Instruments (Castle Hill, Australia).

In the above “acute” protocol, applied for 11 pigs, the animals were monitored for hemodynamic, blood cell and inflammatory mediator changes for 6 h before sacrificing them for organ collection. Blood was collected before and at 5, 10, 15 and 30 min after each injection. In another type of acute studies, the animals obtained the vaccine i.m., once or several times repeatedly. The third type of study was ‘chronic”, i.e., “booster” immunization was applied in the same way as in acute study, except that the animals had previously been treated with intravenous Comirnaty 6 weeks earlier. Blood withdrawals in these pigs were performed at 3 weeks and at 6 weeks, before and during the final “standard” CARPA experiment and organ collection.

### 2.4. Blood Assays

Ten mL of venous blood was drawn from the pigs at different times into EDTA containing vacuum blood collection tubes (K3EDTA Vacuette, Greiner Bio-One Hungary, Mosonmagyaróvár, Hungary). 0.5 mL of blood was aliquoted for use in an ABACUS Junior Vet hematology analyzer (Diatron, Budapest, Hungary) to measure the following parameters of blood cells: white blood cell (WBC), granulocyte (GR) and lymphocyte (LY), platelet (PLT), red blood cell (RBC) count and hemoglobin (Hgb) concentration. For measuring thromboxane B2 (TXB2), a stable metabolite of thromboxane A2 (TXA2), 4 μg indomethacin (diluted in 2 μL of 96% ethanol) was mixed with 2 mL of anticoagulated blood to prevent TXA2 release from WBC before centrifugation at 2000× *g*, for 4 min at 4 ^°^C. Another 2 mL of anticoagulated blood was directly centrifuged using the same settings to separate the plasma.

After centrifugation, the plasma samples were aliquoted, frozen, and stored at −70 ^°^C until the TXB2 assay was performed as described in the kits’ instructions. We used a commercially available ELISA kit (Cayman Chemicals, Ann Arbor, MI, USA) and an FLUOstar Omega microplate reader (BMG Labtech, Ortenberg, Germany).

### 2.5. Isolation of PBMC

The blood samples collected as described above were aliquoted, and one portion was used for peripheral blood mononuclear cell (PBMC) separation within 30 min after blood withdrawal. Briefly, 2 mL blood was transferred into a 15 mL tube and diluted with 2 mL phosphate-buffer saline (PBS pH 7.4). In a new 15 mL tube, 3 mL Ficoll-Paque media (GE Healthcare, Chicago, IL, USA) was pipetted into the bottom and the diluted 4 mL blood sample was carefully layered on top, centrifuged for 30 min at 400× *g*. The upper plasma layer was removed, and the leukocyte layer was transferred into a new tube containing 6 mL PBS, washed and centrifuged.

### 2.6. RT-qPCR analysis of mRNA uptake in PBMC and different pig organs

The PBMCs were resuspended in 1 mL TriZol (Thermo Fisher Scientific, Waltham, MA, USA) and total RNA was extracted according to manufacturer’s instructions. RNA pellet was resuspended in RNAse-free water and the RNA concentration was determined photometrically on a NanoDrop microphotometer (Thermo Fisher). One microgram of RNA of each sample was reverse-transcribed with the high-capacity cDNA reverse transcription kit from Applied Biosystems (Applied Biosystems/Life Technologies, Carlsbad, CA, USA) using random primers in a final volume of 20 μL. Quantitative real-time PCR (RT-qPCR) reactions were then performed on a Bio-Rad CFX96 thermal cycler (Bio-Rad Hungary, Budapest, Hungary) using the SensiFast SYBR Green PCR Master Mix (Thermo Fisher). The specificity and efficiency of each PCR reaction was confirmed with melting curve and standard curve analysis, respectively. Each sample was quantified in duplicate and normalized to the same sample’s 18S rRNA (RN18S) expression. Mean expression values were calculated as fold expression relative to a baseline control sample using the 2^−ΔΔCt^ formula which transforms Ct values into a biologically meaningful fold-change. If fold-change is >1, mRNA is accumulated or the cytokine gene is upregulated, and if it is **<** 1, there is no net mRNA uptake or the gene is downregulated. Pre-designed primers for IP10 (CXCL10, Assay ID: qSscCED0019399) were purchased from BioRad. Primer sequences for SP, TNF-a, RN18S, IL1RA and CCL2 are shown in STable 1.

### 2.7. Histology and immunostaining of pig tissues

Histologic analysis was done by the standard HE staining. For immunostaining, formalin-fixed paraffin embedded kidney, heart and brain tissues were cut and 4 µm slices mounted on SuperFrost Plus slides (Thermo, USA). Heat-induced antigen retrieval was performed with citric buffer pH 6.0 for 20 minutes, then slides were blocked with 5% donkey serum for 30 minutes and stained for SP1 protein (1:400 rabbit monoclonal anti-SP1, Thermo, USA) at 4C overnight. For kidney and heart samples, the secondary antibody (1:200 biotinylated donkey anti-rabbit antibody, Jackson Immunoresearch, USA) was applied for 1 hour followed by HRP-conjugated streptavidine (1:200, Genetex, USA) and developed with Vulcan Fast Red substrate (Biocare, Pacheco, CA, USA)). Immunostaining was evaluated under Leica DM-R light microscope at 200x magnification. Brain sections were incubated with fluorescent secondary antibody (1:200, goat anti-rabbit-Alexa-594, Jackson Immunoresearch, West Grove, PA, USA) for 2 h in the dark. The slides were then covered with coverslips on VectaShield mounting medium containing DAPI (4′,6-diamidino-2-phenylindole) for nuclear staining (Vector Laboratories, Newark, CA, USA) and analyzed under a fluorescent microscope at 400x magnification.

### 2.8. Statistical Analyses

All data are presented as mean ± SD. Statistical analysis was performed using SPSS 10 (IBM, New York, NY, USA). Basic cardiopulmonary parameters were evaluated using paired *t*-test, while blood cell counts and PBMC gene expression values were evaluated using Kruskal–Wallis test and Dunn’s post-hoc test for multiple comparisons. Level of significance was set to *p* < 0.05 in each analysis.

## 3. RESULTS

### 3.1. The acute anaphylactic reactogenicity of Comirnaty in naive and anti-PEG hyperimmune pigs: previous results

Previous studies showed that pigs responded to one-time intravenous administration of Comirnaty with transient hemodynamic, hematologic, and inflammatory mediator changes, most notably pulmonary hypertension, granulocytosis, and thromboxemia; a symptom triad characteristic of complement activation-related pseudoallergy (CARPA). This reaction occurred in 6 out of 14 pigs, and in one case it escalated into anaphylactic shock (Dezsi et al., 2022). The reaction could be substantially aggravated by prior sensitization of the animals through immunization with PEGylated liposomes (Doxebo), which raised the anti-PEG antibody levels by 2-3 orders of magnitude after about 10 days. At that time, we found that all nine hyperimmune pigs developed anaphylaxis to intravenous administration of the human equivalent dose (HED) of Comirnaty (Barta et al., 2024).

### 3.2. Anaphylaxis and concurrent induction of systemic inflammation by Comirnaty

In the present study, we aimed to reproduce the above-mentioned results in naïve and anti-PEG hyperimmune pigs, with a methodological modification. Specifically, to maximize systemic immune activation and facilitate the detection of changes in peripheral inflammatory markers, the immunization phase was intensified by administering three successive injections of Comirnaty, each corresponding to five times (5×) the human equivalent dose (HED). The other steps of anti-PEG immunization and vaccine adminstraion were similoar to the previuos study and the data shown here represent only new and correlating information.

As evidenced by the 200-fold increase in mean anti-PEG IgM levels over baseline one week after Doxebo immunization (Supplementary Figure 1), the pigs were comparably sensitized to Comirnaty-induced anaphylaxis as in our previous study, where all animals developed anaphylactic shock from a single HED of Comirnaty (Barta et al., 2024). However, the cardiovascular responses differed markedly. As shown in Figures 1A and 1B, only 2 of the 10 animals developed anaphylactic shock, with massive changes in pulmonary arterial pressure (PAP), systemic arterial pressure (SAP), and heart rate (HR), requiring resuscitation. In one of these pigs (Figure 1A), the reaction was tachyphylactic, meaning the pulmonary response could not be triggered by the second 5× HED injection. In the other pig (Figure 1B), the life-threatening reaction was non-tachyphylactic.

**FIGURE 1.**
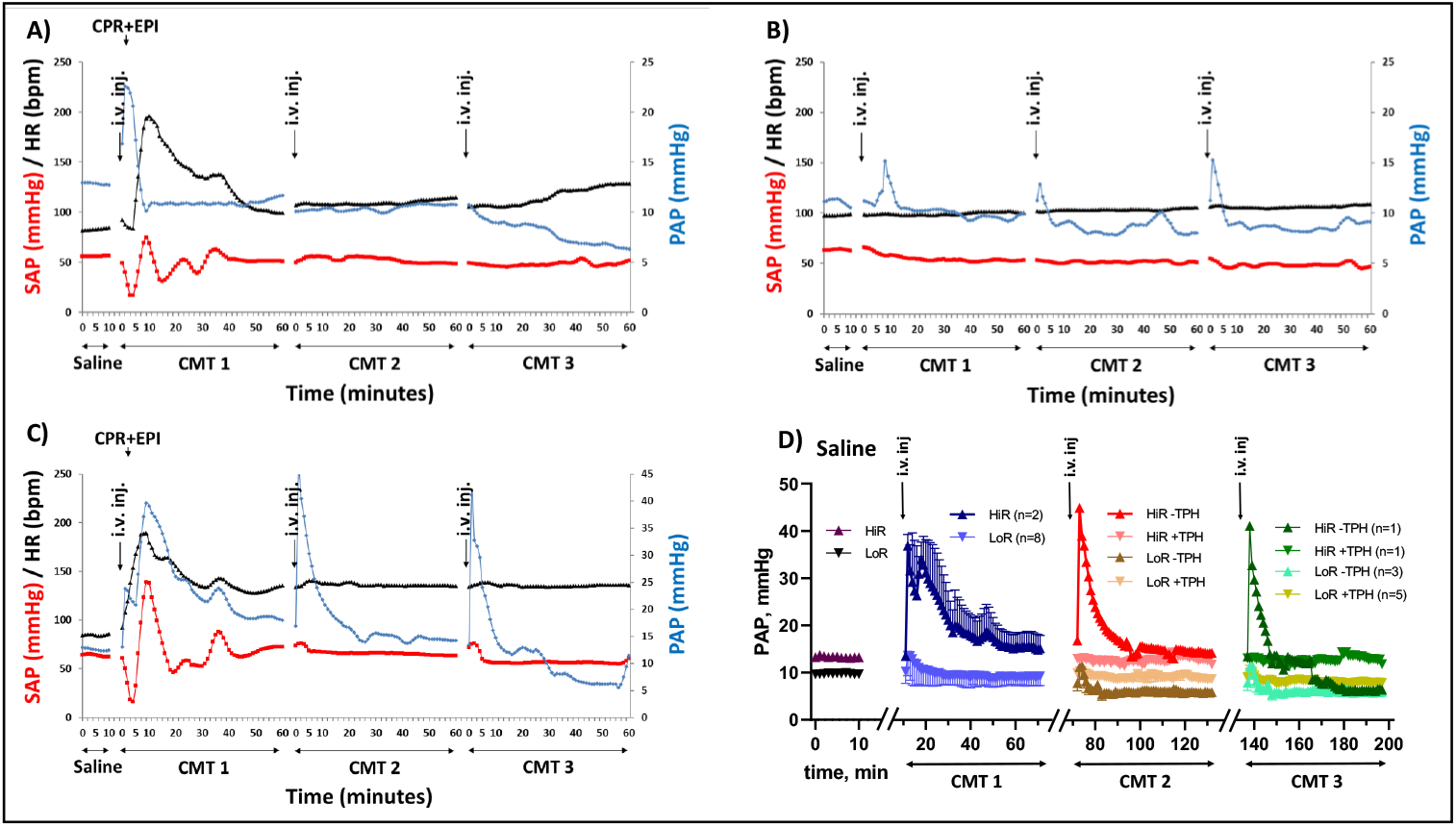
Acute hemodynamic changes after three consecutive injections of Comirnaty (CMT) applied in 5X human vaccine dose, roughly ~ 0.1 mg lipid/kg. A-C. Typical changes out of 10 similar experiments showing the individual variations. SAP – systemic arterial pressure, PAP – pulmonary arterial pressure, HR – heart rate. Three intravenous injections of CMT are denoted by CMT 1 – CMT 2 – CMT 3; CPR+EPI – cardiopulmonary resuscitation with intracardiac administration of epinephrine. D. PAP changes are mean ± SEM and categorized by the type of reactions. HiR – high responders, LoR – low responders; −TPH without, and +TPH - with tachyphylaxis

### 3.3. Blood changes attesting to acute proinflammatory effects of Comirnaty

Figure 2A shows that consecutive administrations of 5X-HED vaccine boluses led to 10-20% leukocytosis after each injection, without return to baseline after the individual injections. These additive rises were most likely due to bone marrow release of stored neutrophils and led to a lasting (> 6 h), up to 60% leukocytosis, implying active inflammatory, infectious, or immunological process (Christopher & Brounts, 2010; Ndeupen et al., 2021). The proinflammatory nature of mRNA-LNPs have been associated with a vicious cycle of cytokine release and complement (C) activation (Ndeupen et al., 2021; Nurnberger et al., 1994; Szebeni, 2025; Szebeni, 2025; Szebeni & Koller, 2025), and the data presented in Figure 2 show parallel induction of these processes as well, thus proving that the leukocytosis is a consequence of the vaccine’s proinflammatory effect.

**FIGURE 2.**
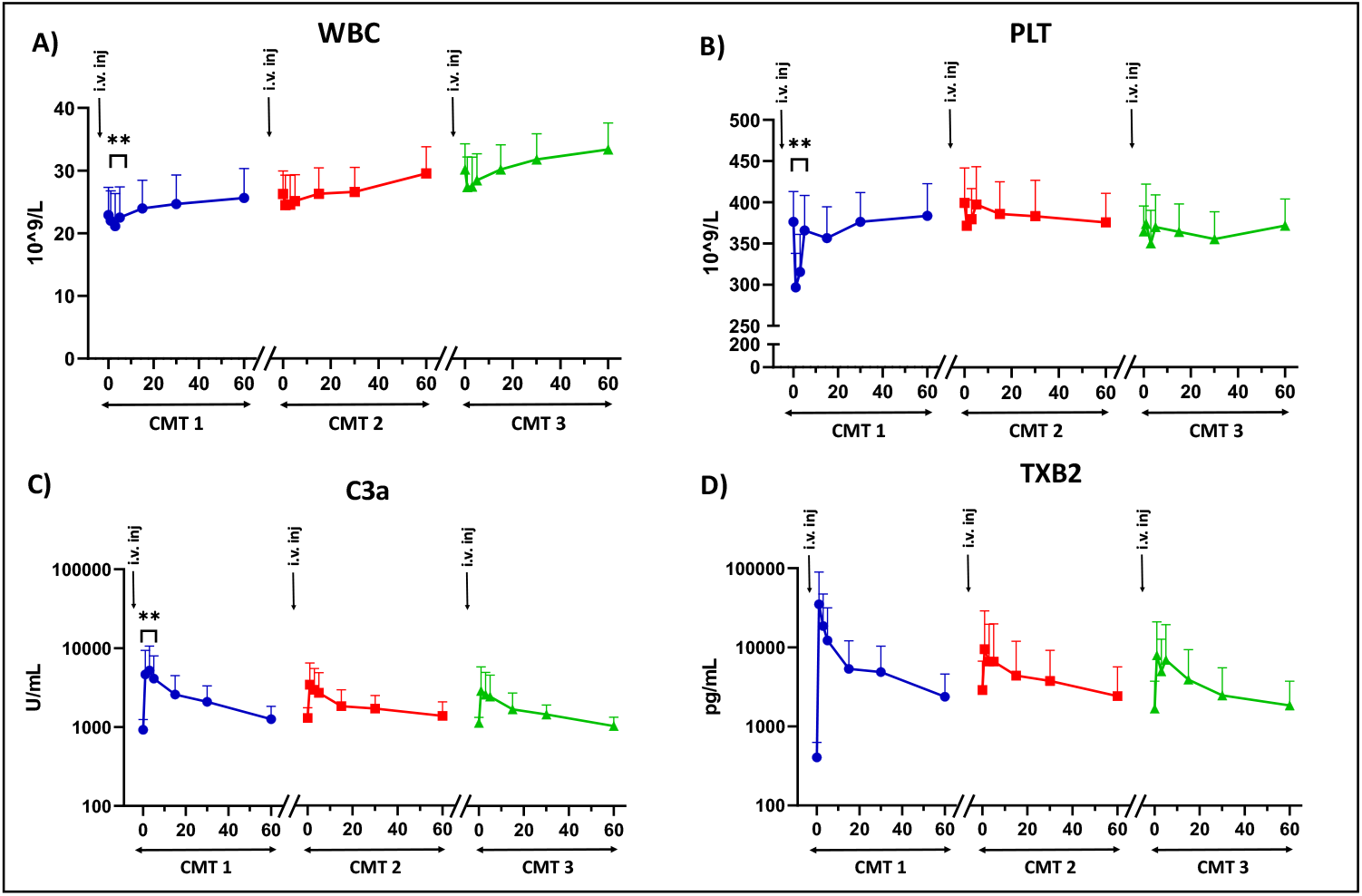
Acute changes in white blood cell (A) and platelet (B) C3a (C) and TXB2 (D) counts after three consecutive injections of Comirnaty in pigs. Mean ± SEM, n= 9 pigs

Figure 2B shows the other characteristic blood cell effect of immune active nanoparticles: transient thrombocytopenia. This change was observed only after the first vaccine injection, and it could not be repeated with further injections. Such thrombocytopenia is a well-documented consequence of leukocyte and platelet activation, as well as pulmonary endothelial cell stimulation mediated by the anaphylatoxins C3a and C5a (Szebeni & Koller, 2025). This process involves the formation of platelet-leukocyte aggregates and their transient adhesion to pulmonary endothelium, a phenomenon referred to as pulmonary marginalization. These interactions contribute to platelet sequestration and reversible thrombocytopenia during systemic inflammatory reactions (Guo & Ward, 2005; Sims & Wiedmer, 1991).

Consistent with the above-delineated role of C activation in leukocytosis and thrombocytopenia, we found laboratory evidence of C activation: sudden rise of C3a anaphylatoxin on the same time course where blood cell changes occur (Figure 2C). Similar changes were observed with TXB2, the stable metabolite of TXA2, the immediate effector of pulmonary and coronary vasoconstriction (Figure 2D).

The reaction was partially tachyphylactic, just as the hemodynamic response in some animals, attesting to causal relationship between these processes (Bakos et al., 2024).

### 3.4. Pilot study findings on intramuscular vaccine-induced CARPA

To address concerns regarding the clinical relevance and translatability of the pig experiments described above - specifically the use of intravenous versus i.m. vaccination - we conducted a preliminary investigation to assess whether i.m. administered vaccine boluses could induce CARPA. The study involved i.m. injection of two pigs with 5X-HED of Comirnaty: one developed tachycardia and fever, while the other, which experienced rebleeding at the injection site, exhibited signs of anaphylaxis. These findings suggest that even i.m. vaccination may result in systemic exposure through inadvertent intravascular injection, lymphatic drainage, or inflammation-induced increases in vascular permeability (J. Szebeni & Koller, 2025). Nonetheless, given the limited sample size, no conclusions can be drawn regarding the incidence or predictability of such reactions after i.m. administration of the vaccine, which supports the use of intravenous delivery in the present experiments.

### 3.5. Uptake of mRNA-LNP

#### 3.5.1. mRNA uptake by peripheral blood mononuclear cells (PBMC)

As shown in Figure 3, three consecutive injections of Comirnaty led to variable SP mRNA baseline within an hour. Maximum mRNA uptake, corresponding to 400–600-fold increases over baseline, was observed in two pigs out of 9 (#2 and #7), while the remaining animals showed either clearly detectable (#1 and #3) or minimal increases (#4, #5, #6, #8, and #9). Nevertheless, uptake up to about 100-fold increase over baseline was observed with most animals after 6 h. The insert in Figure 3 further demonstrates that in the chronic study, a similar biphasic rise and fall of SP mRNA levels occurred following injection of the same vaccine doses six weeks after the initial treatment, which emulates booster vaccination. This indicates that neither the initial acute anaphylactic reaction and later, the gradually developing anti-SP and anti-PEG immune response interfered with the instant mRNA-LNP uptake of PBMC. accumulation in PBMCs across different pigs, peaking around 15 minutes and returning to near

**FIGURE 3.**
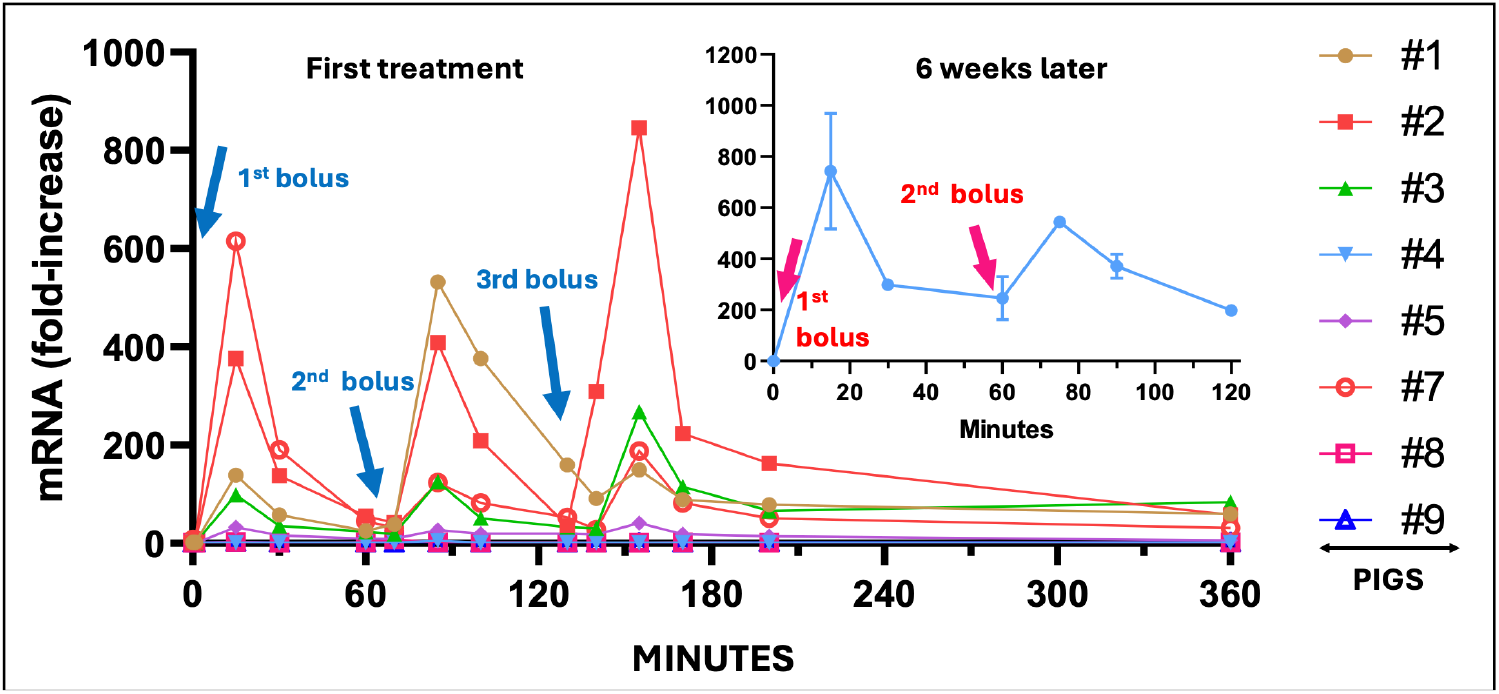
Kinetics of SP mRNA uptake by PBMCs of pigs following three consecutive bolus injections of Comirnaty (blue arrows). Fold increase means the mRNA level normalized to baseline in SPF animals. Curves with different symbols are individual data in 9 pigs, labeled with numbers (see key). The insert shows corresponding data six weeks after the initial Comirnaty treatment, simulating the effects of booster immunization. Here the injections are shown with red arrows.

The human relevance of these findings is noteworthy. As detailed in the Discussion, several studies have shown that a small fraction (~3%) of mRNA-LNPs can enter the circulation shortly after i.m. injection. thje present pig data indicate that this fraction can be rapidly taken up by blood mononuclear cells, initiating inflammatory activation and SP production. The reproducible, short-latency uptake followed by rapid decay, even at 6 hours, was unexpected given the extended efforts to stabilize the mRNA via chemical modifications (Kariko et al., 2005; Kauffman et al., 2016; Pardi et al., 2018; Schlake et al., 2012). These results suggest a non-saturable uptake mechanism that, despite rapid consumption and/or metabolism, efficiently induces cytokine expression. Together, they provide the first direct evidence supporting the pig as a suitable model for studying multisystem inflammatory syndrome.

#### 3.5.2. Uptake of mRNA by pig organs

Figure 4 shows that 6 h after vaccine injections, in addition to PBMC, 6 other organs of 9-11 pigs displayed varying increases of cellular levels of SP mRNA, indicating universal, although individually highly variable transfection of organ cells with SP mRNA. For example, the livers of all 11 pigs displayed SP mRNA accumulation, with variance over 3 orders of magnitude. The splenic uptake in 9/11 animals ranged between 10 and 3000-fold, while the corresponding lymph node and PBMC ratios were 10/11 and 7/11, respectively, up to about 100-fold increase. The kidney and heart uptake in 7/11 and 4/11 pigs ranged up to 10-fold, while near all, 9 of 11 pigs showed a minimal uptake of mRNA by the brain. These data provide evidence that the vaccine nanoparticles can transfect both the blood cells and different pig organs on a time scale of minutes to hours after entering in blood.

**FIGURE 4.**
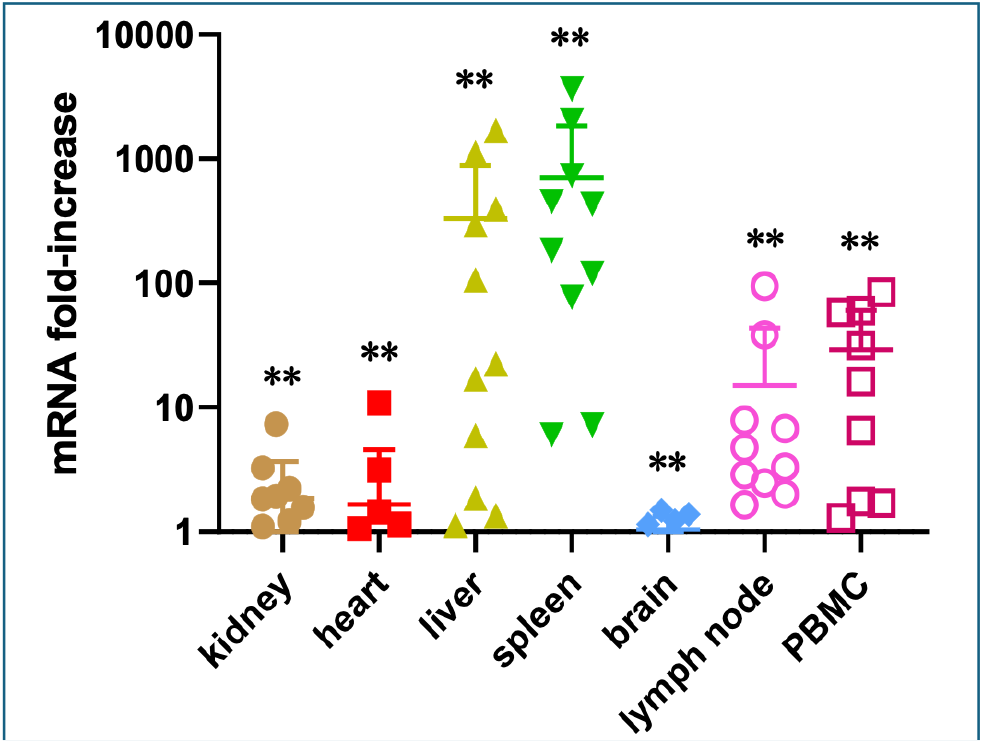
Organ transfection with SP mRNA in pigs 6 h after injection of Comirnaty. Values are >1 fold-increases (mean ± SD, n=9-11) from the RT-qPCR analysis relative to the SPF control. Values < 1 were considered 0 (baseline). Different symbols visualize individual values in different organs.

### 3.6. Upregulation of proinflammatory cytokine genes in PBMC

Importantly, the transient accumulation and rapid degradation cycles of SP mRNA in PBMC (Figure 3) were associated with upregulation of proinflammatory cytokine genes IL1-RA, CXCL10, TNF-7 and CCL2 at 6 h post-injection (Figure 5), indicating inflammation in blood. Thus, we detected signs of systemic inflammation in the model, which is the essence of MIS. Moreover, the extents of different cytokine gene upregulations substantially varied among the animals, which is consistent with the unpredictable rare occurrence of MIS in humans. To mention among other differences, the CCL2 gene upregulation in the 5-60-fold range was significantly more expressed than the maximal 10-fold increases observed for the other 3 cytokines’ mRNA, although IL-1RA was particularly expressed in the PBMC of 2 of 11 pigs. In keeping with the Toll-like receptor 4-mediated activation of transcription factors NFkB and interferon regulatory factor (IRF) in a monocyte cell line (Zelkoski et al., 2025), these cytokine releases can most easily be explained by monocyte activation.

**FIGURE 5.**
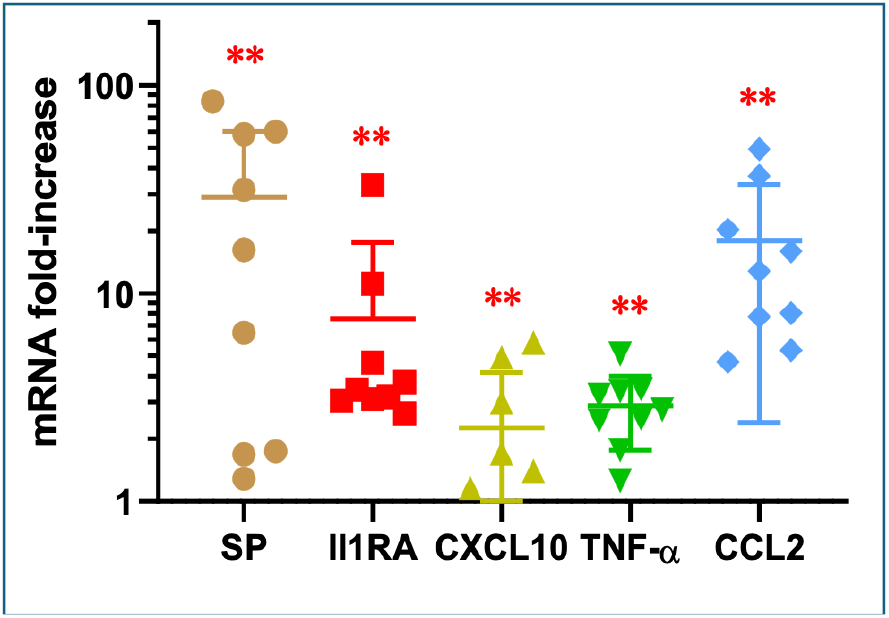
Upregulation of proinflammatory cytokine genes in PBMC 6 h after injection of Comirnaty (mean ± SD, n=9). Different symbols visualize different genes in individual animals. **, P, 0.05.

### 3.7. Upregulation of proinflammatory cytokines in pig organs

Similarly to PBMC, the vaccine-induced SP mRNA was associated with variable upregulation of different proinflammatory genes in different pig organs at 6 h after injection of Comirnaty. As shown if Figure 6, the liver accumulated the most SP mRNA, which was followed by up to 100-fold accumulation of IL1-RA, and up to 10-fold CXCL10, TNF-7 and CCL2 genes at 6 h post-injection

**FIGURE 6.**
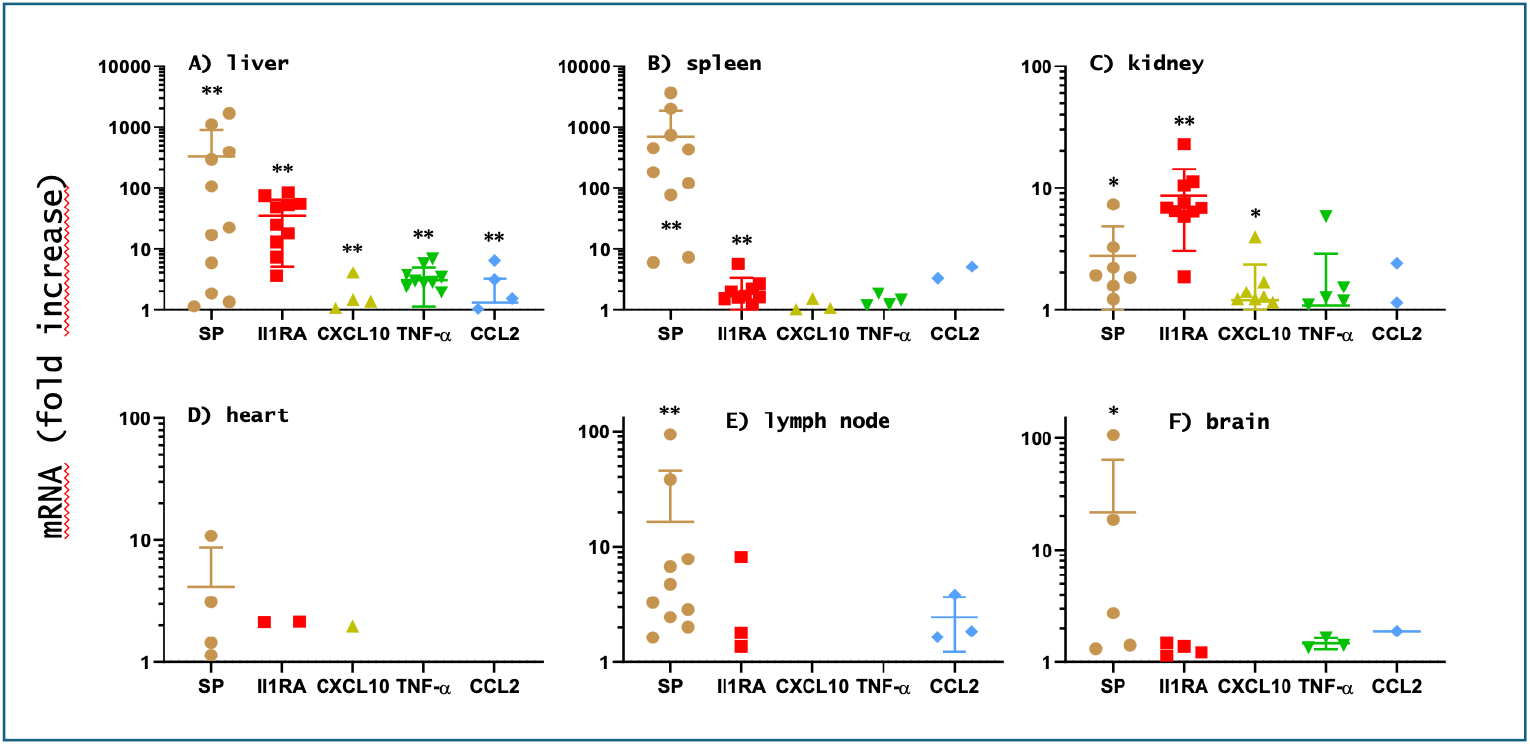
Association of SP mRNA transfection with proinflammatory cytokine gene upregulation in different pig organs 6 h after Comirnaty injection (mean ± SD, n=9-11). Various symbols visualize differences in gene upregulation. Symbols for each gene identify different animals.

Overall, these data prove the utility of the model for quantitative assessment of the organ distribution of intact SP mRNA after vaccination and the consequent systemic inflammatory status of the animals. In other words, the experiment provide a model for multisystem inflammatory syndrome.

### 3.8. Histologic alterations in the different in pig organs

We found no significant pathological changes in sections of skin, lymph node, myocardium, and spleen tissues 6 hours after vaccination. However, as shown in Figure 7, we could observe neuropil vacuolization in almost all brain samples and vascular edema in some cases. Unless artifacts during tissue processing, these are histopathological findings in the dense network of axons, dendrites, and glial processes, and may indicate some kind of tissue damage, such as hypoxic-ischemic injury. Livers of pig #1 and #9 showed mild sinusoidal hyperemia, and in the case of pig #3, we detected multifocal necrosis and moderate periportal hepatitis. Kidneys of #4, #5, and #9 showed multifocal tubular dilation. Tubular epithelial cells were either flattened or cuboidal. STable 2 details all 6h histopathological data from 7 organs in 8 pigs.

**FIGURE 7.**
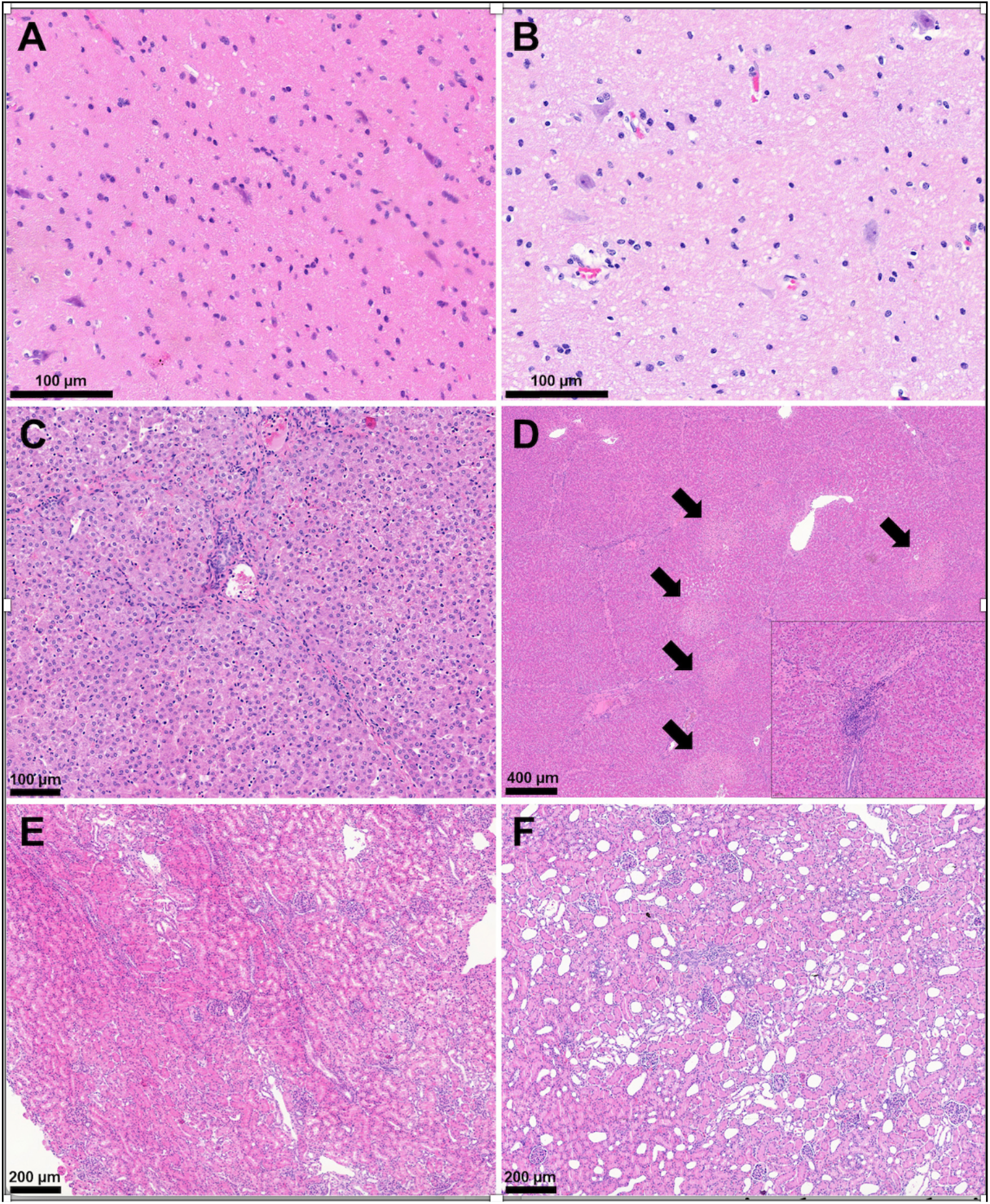
Comparison panel of SPF control tissues (A, C, E) and specimens of vaccinated pigs containing pathologic lesions (B, D, F). A: brain tissue with normal histologic structure; B: brain tissue showing mild vacuolization of neuropils, and vascular edema (bar=50 µm). C: normal architecture of liver; D: multifocal necrosis in the liver parenchyma (arrows). The inset shows the moderate mononuclear cell infiltrate at the portal/periportal area (bar=500 µm, inset: bar=100 µm). E: renal tissue with physiologic histologic structure; F: multifocally and mildly dilated tubules in the cortex of the kidney (bar=200 µm). All sections are stained with H&E.

### 3.9. SP protein expression in pig organs

Immunostaining for SP in kidney and heart tissues revealed a strong positive reaction in renal tubules (Figure 8A) and a milder positivity in the myocardium (Figure 8B) of vaccinated pigs (right panel), as compared to the tissues of non-vaccinated control (left panel). SP protein was also expressed in brain cells (Figure 8C, see arrows) of vaccinated pigs 6 h after the injection of Comirnaty (lower panel), but was not detected in non-vaccinated pig brain tissue.

**FIGURE 8.**
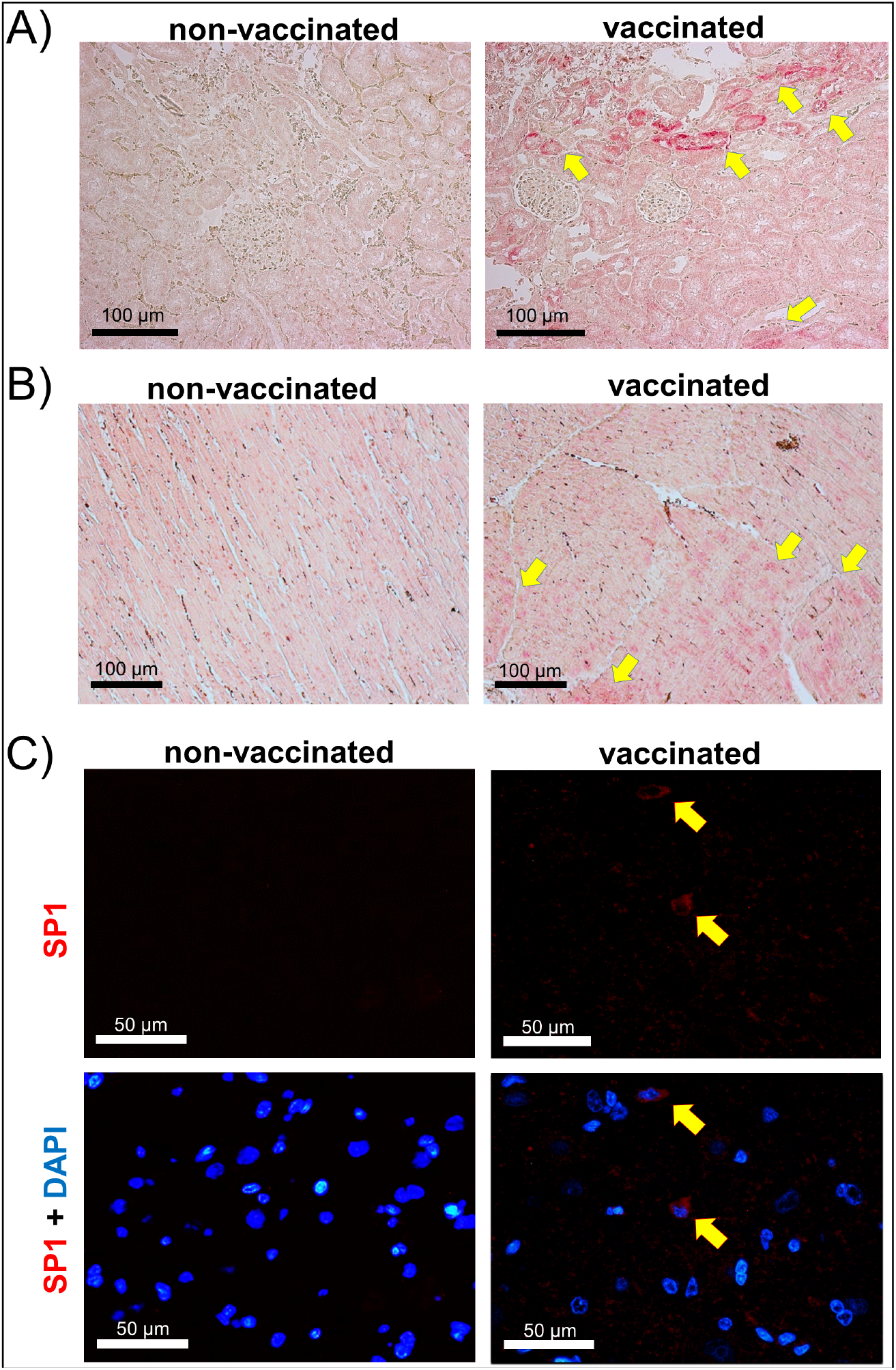
Immunohistochemistry for SP protein expression in pig kidney (A), heart (B) and brain (C) 6 h after injection of Comirnaty or vehicle. Arrows point on the SP1-positive stain of kidney tubules (A) and myocardial tissue (B) as well as SP1-positive brain cells (C). Scale bar=100µm (A, B) and 50µm (C). DAPI (blue) stains the nuclei in immunofluorescent slides.

## 4. DISCUSSION

### 4.1. The challenge of vaccine-induced adverse events

Since the introduction of mRNA-based COVID-19 vaccines designed to encode antigens instead of delivering them directly, concerns are growing about a broad spectrum and relatively high incidence of severe AEs. The phenomenon has led to the distinction of a novel diagnostic category: “post-vaccination syndrome” (PVS) (Bhattacharjee et al., 2025; Krumholz et al., 2023; Oueijan et al., 2022; Palmer et al., 2023; Shrestha & Venkataraman, 2024), whose chronic, occasionally disabling complications can be considered as an iatrogenic orphan disease (Szebeni, 2025). While the absolute incidence of all-type AEs is low (0.03–0.5%) (Szebeni, 2025), the global administration of multibillion doses implies that the number of PVS cases is in the millions. The seriousness of the issue is underscored by the fact that several countries have suspended, restricted, or issued contraindications for mRNA COVID-19 vaccines. In the US, legislative initiatives to limit or ban these vaccines have been introduced in multiple states. As of May 2025, the US health authorities revised their recommendations to prioritize mRNA vaccination for individuals at higher risk for severe illness. Since then, the reassessment of vaccine safety and efficacy, along with necessary changes in vaccination policies, has continued to evolve, making rigorous scientific investigation into this divisive medical issue essential.

### 4.2. Study goals and Rationale

The goal of this study was to investigate potential links between intravenous administration of Comirnaty, cardiopulmonary hemodynamic changes, multiorgan mRNA transfection and inflammatory cytokine gene upregulation in pig blood and organs. Prior immunization of the pigs with PEGylated liposomes sensitized the animals to the PEGylated LNP carrier of the vaccine and thus accelerated the acute innate response manifested in anaphylaxis. It was hypothesized that simultaneous monitoring of anaphylaxis endpoints, mRNA uptake in blood and organs, their translation to SP and cytokine gene upregulation reproduces key features of human vaccine-induced anaphylaxis and multisystem inflammatory syndrome in a single step, providing a translational value to the model.

This setup deviated from standard human COVID-19 vaccination practice in a major respect: it involved intravenous administration of a 15-fold higher vaccine dose compared with the human intramuscular protocol. The rationale for this modification, as well as for using pigs hypersensitized to anaphylaxis, was that the mechanisms of rare diseases, such as vaccine-induced anaphylaxis and multisystem inflammatory syndrome, can only be studied in amplified animal models where the frequency of disease markers is sufficiently high for statistical analysis. In our case, reproducing the human incidence rates of anaphylaxis and multisystem inflammatory syndrome would have required an impractically large number of animals, corresponding to the human non-responder/responder ratio in the 10^4^-10^6^ order for each positive case.

An amplified model of anaphylaxis was recently developed by modifying the naturally hypersensitive porcine CARPA model (Szebeni & Bawa, 2020; Szebeni, 2024). In this hypersensitivity model intravenous administration of a human-equivalent dose of Comirnaty caused anaphylactic shock in all nine pigs previously vaccinated with PEGylated liposomes (Doxebo), which had raised their anti-PEG antibody levels by several orders of magnitude (Barta et al., 2024). In this model, the elevation of the anaphylatoxin C3a, associated with the anaphylactic shock, indicated a causal role of complement activation in the reactions (Barta et al., 2024). In this study, C3a was measured to explore the possible role of systemic complement activation in organ inflammation, since complement is a primordial driver of inflammation in general, including long-COVID (Cervia-Hasler et al., 2024). However, individual variations in the analytes prevented establishing a significant correlation between C3a elevation and cytokine gene upregulation (data not shown).

The cytokines investigated are critically involved in mediating inflammation and immune cell recruitment during immune responses and pathological conditions (Jaffer et al., 2010; Kany et al., 2019). Among them, IL-1 receptor antagonist (IL-1RA), CXCL10, tumor necrosis factor-alpha (TNF-α), and CCL2 play distinct yet interconnected roles. IL-1RA is an anti-inflammatory cytokine that inhibits IL-1 signaling, often upregulated as a compensatory response during inflammation. CXCL10 is a chemokine that recruits Th1 cells and NK cells, commonly elevated in viral and autoimmune conditions. TNF-α is a central proinflammatory cytokine that promotes cell recruitment, apoptosis, and systemic inflammation. CCL2 attracts monocytes and contributes to chronic inflammation and tissue damage. Together, these mediators reflect and drive inflammatory responses across diverse pathological settings (Jaffer et al., 2010; Kany et al., 2019).

As mentioned, intravenous delivery of Comirnaty was used to ensure mRNA-LNP exposure to blood, which is in keeping with evidence of vaccine entry into the bloodstream following i.m. administration in humans (Borresen et al., 2020; Ltd, 2021; Ndeupen et al., 2021). In fact, rapid entry of i.m.-injected LNPs into the bloodstream had been described over a decade before the use of LNP for vaccination (Pardi et al., 2015), and a preclinical rat study on Comirnaty pharmacokinetics also showed leakage from the injection site and widespread distribution of labeled LNPs throughout the body, starting within minutes (Pfizer Australia Pty Ltd, 2021). The latter experiment showed ~3% of injected LNP lipid in blood within 15 min after i.m. injection, and that its’ plasma level doubled within 2-6 h (Pfizer Australia Pty Ltd, 2021). As to the journey of the vaccine from the deltoid muscle into blood, a recent review (Szebeni & Koller, 2025) discussed 3 options: (i) accidental injection into a blood vessel, (ii) natural lymphatic drainage and (iii) reversed EPR effect (Enhanced Permeability and Retention), i.e., enhanced tissue to blood permeation of nanoparticles along their concentration gradient due to inflammation-related permeability increase (Azzopardi et al., 2013).

### 4.3. Basic observations and novel findings

The reproduction in the present study of Comirnaty-induced peudoallergy and anaphylaxis in pigs that was associated with C3a and thromboxan B2 elevations strengthene the CARPA concepts explaining these reactions. However, as detailed below, some observations were unexpected and represent new information on this phenomenon.

The highly variable, but in some animal robust uptake of functional SP mRNA by blood cells and main organs implies multisystem transfection of functional mRNA, a well-known feature of mRNA-LNPs leading to their clinical development as nucleic acid delivery agents (Cheng et al., 2023; Cullis & Felgner, 2024; Francia et al., 2020). As expected from the tissue distribution of mRNA-LNP reported in the literature, the liver, spleen and lymph nodes showed the most uptake in the overwhelming number of pigs, as well as the PBMC, which too contains LNP target cells (monocytes). What was a less predictable result is that the heart and the kidney of a portion of animals also took up the mRNA, which are not target tissues for the vaccine, yet display AEs, particularly the heart. Regarding the mechanism of mRNA-LNP uptake by non-phagocytic cells, it can be through endocytosis, either clathrin or caveolae mediated or micropinocytosis. This process is enhanced in inflammation**-**activated endothelium (J. Szebeni & Koller, 2025). The capability of LNPs for fusion, due to their ionizable lipid-content, proceeds only in acidic medium and is implicated in the endosomal escape of mRNA-LNPs (Cheng et al., 2023; Cullis & Felgner, 2024; Francia et al., 2020). Whether or not it plays a role in multiorgan transfection, is unclear.

The present study led to several original observations. Regarding anaphylaxis, our findings indicate that the pro-anaphylactic effect of high anti-PEG antibody level in blood is neither obligatory nor universal. Notably, in the present study, a vaccine dose 15-fold higher than that in our previous report (Barta et al., 2024) resulted in more expressed individual variation of reaction strength and the presence or absence of tachyphylaxis, entailing lower incidence of anaphylactic shock. Although the reason of this inter-experimental variation has not been clarified, a biphasic dose-effect relationship cannot be excluded. Such nonlinear response patterns are well known in immunology, exemplified by the immune-stimulating and suppressive effects of low- and high doses of lipopolysaccharide via TLR4 pathway modulation (Lajqi et al., 2021; Lajqi et al., 2019); or the IL-2 signaling in T cells (Boyman & Sprent, 2012) and IFN-γ stimulation of macrophages (Schroder et al., 2004).

The novelties in our SP expression and cytokine upregulation data, taken as indicators of mRNA transfection and organ inflammation, include a temporal, but not quantitative, association between mRNA uptake and the upregulation of inflammatory cytokines across different tissues. In PBMC, the identity of WBC subtypes responsible for mRNA uptake remains unclear. Considering that in PBMC, consisting of T cells (~60–80%), B cells (~5–15%), NK cells (~5–15%), and monocytes (~10–20%), only monocytes have the capability for endocytic and phagocytic uptake of mRNA-LNPs (Ndeupen et al., 2021; Pardi et al., 2018; Wei et al., 2024), monocytes are the most likely candidates for mRNA accumulation and cytokine gene upregulation. This proposal is in keeping with the fact that mRNA-LNPs were shown to activate monocytes through Toll-like receptor 4 (TLR4), MyD88-dependent signaling and NLRP3 inflammasome activation, leading to the release of proinflammatory cytokines and chemokines into blood (Ndeupen et al., 2021; Pardi et al., 2018; Wei et al., 2024). Since these processes in monocytes are associated with lysosomal disruption and the release of cathepsins into the cytosol, the entailing apoptosis (Zelkoski et al., 2025) explains the vaccine’s toxicity on monocytes (Bhattacharjee et al., 2025).

Regarding the organ uptake of mRNA and cytokine upregulation, it showed relatively high reproducibility of transcription profiles among animals, contrasted by substantial variation among tissues, adding a tissue-specific dimension to the transfection/inflammatory response. We also found evidence of SP protein appearing in the kidney and heart 6 h after vaccination, supporting rapid translation of the mRNA into protein. However, because of this rapid translation of SP mRNA to SP, the individual roles of the mRNA, the LNP and the SP in the driving of inflammation remains unclear.

### 4.4. Clinical perspective on two cardinal symptoms of multisystem inflammatory syndrome: gut and heart inflammations

In addition to confirming the CARPA concept of anaphylaxis and extending current knowledge of this innate immune reaction to mRNA vaccines in humans, the minute-by-minute alignment between vaccine mRNA transfection and the onset of inflammatory cytokine upregulation in vaccine-vulnerable organs provides strong evidence for a causal link between these phenomena. This represents an important step toward elucidating the mechanisms underlying not only multisystem inflammatory syndrome, but also its single-organ manifestations, among which gastrointestinal and cardiac inflammations are among the most frequent (Elsaid et al., 2023; Feldstein et al., 2020; Hoste et al., 2021; Ruvinsky et al., 2022; Whittaker et al., 2020).

Acute vaccine-induced gastroenteritis has been hypothesized to be a secondary consequence of prior manifest or latent COVID-19 infection (Hoste et al., 2021; Ruvinsky et al., 2022), supporting the concept of the vaccine acting as a “second inflammatory hit” (Hoste et al., 2021; Ruvinsky et al., 2022). In this context, recent findings on COVID-19–induced gut microbiome pathology (Brogna et al., 2022; Brogna et al., 2023; Brogna et al., 2024) may hold relevance for both the prevention and treatment of vaccine-induced gastroenteritis.

Because of the life-threating potential of vaccine-induced pericarditis and myocarditis (i.e., myopericarditis), this AE has received the most attention in the AE context (Bozkurt et al., 2021; McCullough & Hulscher, 2025; Sindet-Pedersen et al., 2023). Numerous studies have identified major pathological alterations in the heart, including the presence of biochemically modified SP mRNA and its +1 ribosomal frameshifted peptide products in individuals with carditis - persisting up to six months post-vaccination (Boros et al., 2024)-, as well as SP subunit monomers in vaccine-exposed cardiomyocyte cultures, capable of intracellular aggregation into covalently bonded high-molecular-weight complexes (Schreckenberg et al., 2024; Schreckenberg et al., 2025). These pathological features have been linked to functional anomalies, thereby providing insights into the mechanisms underlying the cardiac AEs. Our data indicate that direct cardiac transfection and inflammation can occur as early as six hours after vaccine administration, suggesting that pathological changes may begin shortly after vaccination, even though the clinical manifestations of AEs often emerge only a long time after primary or booster vaccinations.

### 4.5. Integrated hypothesis on the mechanism of vaccine-induced multiorgan inflammation

Considering the non-organ specific pathology of acute multiorgan inflammatory syndrome, the pathogenesis must involve a tissue element common to all organs. As reviewed recently (J. Szebeni & Koller, 2025), the endothelial cell layer of capillaries is the primary candidate for this common structural element, accounting for ~80–90% of the total blood vessel surface area in the human body (Goncharov et al., 2020). It has therefore been hypothesized that endothelial cells represent the frontline of systemic encounter with mRNA-LNPs, fueling the progression of multiorgan inflammation. Schematic Figure 9 depicts the cascade of cellular and molecular interactions underlying endothelitis and the subsequent development of organ disease.

**FIGURE 9.**
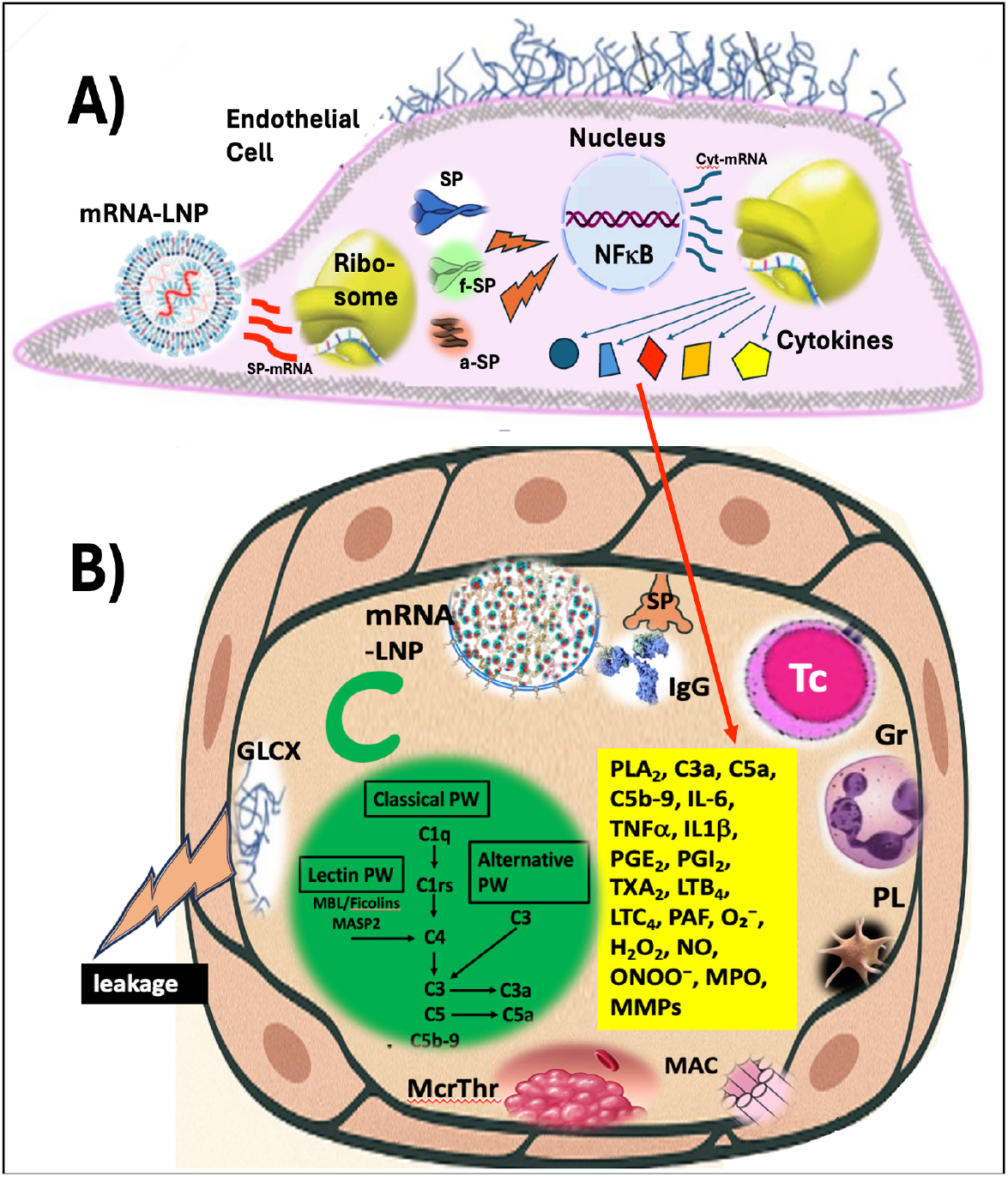
Hypothetic cascade of cellular and molecular interactions underlying the adverse immune activation and inflammatory tissue injury triggered by mRNA–LNP vaccines. **(A)** Schematic representation of endothelial cell (EC) transfection with SP mRNA delivered by mRNA– LNPs. After endocytic uptake and endolysosomal escape, the SP mRNA is translated by ribosomes into full-length SP and aberrant byproducts, including frameshifted (fSP) and aggregated (aSP) forms. These products exert toxic effects on ECs, involving NF-κB–mediated nuclear activation and release of proinflammatory cytokines into the capillary blood. **(B)** Capillary-level mechanisms of vaccine-induced organ inflammation. Pathogenic triggers include direct mRNA–LNP binding to ECs, complement (C) activation via anti-PEG and anti-SP IgG, and additional C activation by SP expressed on the EC membrane. Further processes include cytotoxic T cell (Tc)–mediated attack, adhesion of leukocytes (neutrophils, monocytes, macrophages) and platelets to ECs; cell injury caused by the membrane attack complex (MAC), microthrombus (McrThr) formation, glycocalyx (GLCX) degradation, and capillary leakage. The yellow box highlights key inflammatory mediators contributing to these effects, in addition to the cytokines measured in this study. Tc, cytotoxic T cell; Gr, granulocytes; O_2_−, superoxide; H_2_O_2_, hydrogen peroxide; NO, nitric oxide; ONOO−, peroxynitrite. Panel B was reproduced from Ref. (J. Szebeni & Koller, 2025).

### 4.6. Study limitations and outlook

The deviations of our vaccine treatment protocol from the standard human dose and i.m. administration route were justified by the need for using amplified models to study rare diseases. Among other limitations, the small number of animals was in accordance with the Reduction and Refinement elements of the “3Rs” principle in animal experimentation. Despite the limited dataset for each variable, the results were sufficiently robust to support clinically relevant conclusions. The sole aim of this study was to provide rigorous, evidence-based mechanistic insights into the acute inflammatory adverse effects of mRNA vaccines, without taking position on the broader, polarized debate over these vaccines’ benefits and risks. A better understanding and mitigation of these AEs will contribute to making the mRNA-LNP technology safer.

## CRediT authorship contribution statement

Conceptualization, JSz, GK, LD, methodology and formal analysis GK, GSz, LD; writing/editing: JSz, GK, GSz, LD; investigation: LD, GK, CsR, TM, RF, ASz, TB, GTK, ABD; project administration and funding, JSz, TR, LD and MB. All authors have read and agreed to the published version of the manuscript.

## Institutional Review Board Statement

The study was conducted in accordance with the Declaration of Helsinki, and the animal study protocol was approved by the Institutional and National Review Board/Ethics Committees, PE/EA/843-7/2020.

## Data Availability Statement

Experimental data are available upon reasonable request to the corresponding author.

## Acknowledgments

The expert technical support by Dóra Szkrajcsics, Maria H. Velkei, Katalin Simay, Henriett Biró and Krisztina Fazekas are gratefully acknowledged.

## SUPPLEMENTARY information

**STable 1.**
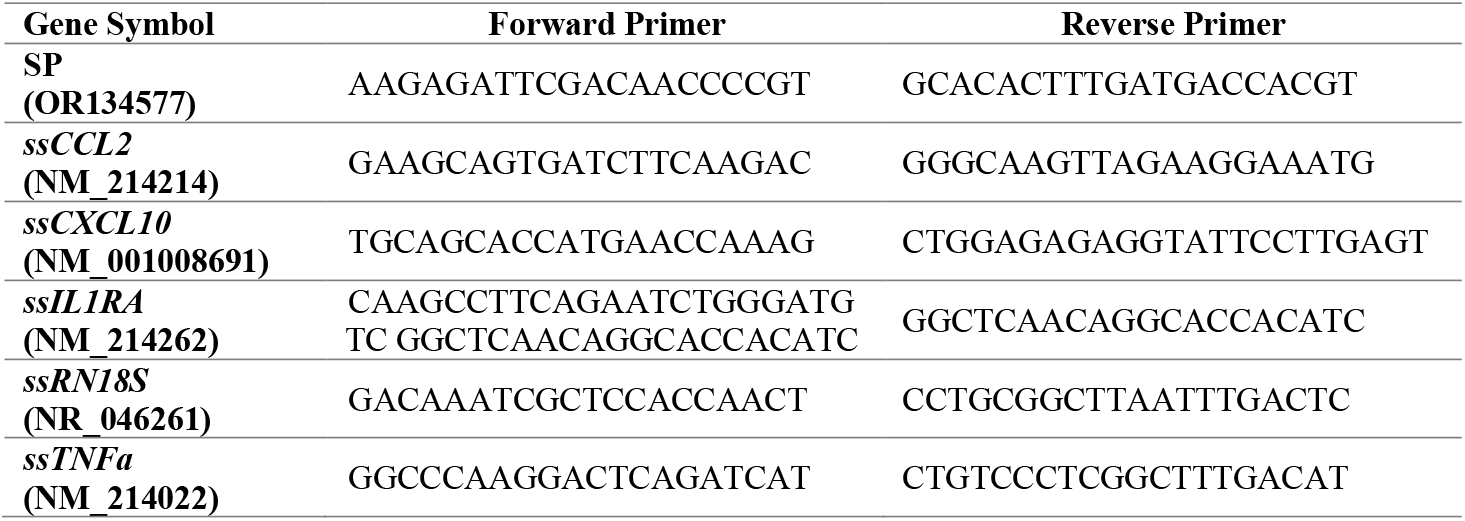
Primer sequences for qPCR analysis.

**STable 2.** Histopathological evaluation of organ specimens obtained from pigs 6h post-treatment.

| Tissue type | Skin | Brain | Cardiac muscle | Lymph node | Spleen | Liver | Kidney |
| --- | --- | --- | --- | --- | --- | --- | --- |
| Code |  |  |  |  |  |  |  |
| #16 | no lesions | no lesions | no lesions | well recognizable follicular structure | well recognizable follicular structure, passive hyperemia due to anesthesia | physiological prominent interlobular connective tissue septae, normal histological structure, intact portal tracts, mild intracytoplasmic vacuolization diffusely | intact glomerular and tubular structure |
| #1 | no lesions | NA | no lesions | mild lymphopenia | no lesions | mild sinusoidal and perivascular hyperemia | no lesions |
| #2 | NA | NA | no lesions | no lesions | no lesions | no lesions | no lesions |
| #3 | no lesions | mild neuropil vacuolisation | no lesions | NA | no lesions | multifocal hemorrhages and single cell to massive necrosis; focal, mild neutrophilic and lymphocytic periportal infiltration; moderate intracytoplasmic vacuolisation diffusely | mild basilar intracytoplasmic vacuolisation in the tubular epithelial cells |
| #4 | NA | mild neuropil vacuolisation | no lesions | no lesions | no lesions | no lesions | multifocal tubular dilation with flattened tubular epithelial cells |
| #5 | no lesions | mild neuropil vacuolisation | no lesions | mild lymphopenia | no lesions | no lesions | multifocal prominent tubular dilation with normal cuboidal tubular epithelial cells |
| #8 | no lesions | mild vascular edema; mild neuropil vacuolisation | no lesions | no lesions | no lesions | no lesions | no lesions |
| #9 | no lesions | mild vascular edema; mild neuropil vacuolisation | NA | NA | no lesions | mild sinusoidal and perivascular hyperemia (v. centralis) | multifocal tubular dilation with flattened tubular epithelial cells |

Pig with code #16 is a control animal with physiological histology of organs (SPF animal). NA: not applicable/no organs were submitted to evaluation. STable 2 shows mild neuropil vacuolisation of the brain specimens in almost all cases, and mild vascular edema in #8 and #9 – this can correlate with hypoxia-induced edema. Still, it can also appear artifactually due to processing. However, we could not detect neuropil vacuolisation, nor vascular edema in the control animal’s brain tissue, therefore, the possibility of being only an artefact is low. Skin samples were taken from the administration sites. Still, no pathologic lesions were detected. Although there was mild reactivity with immunostaining of SP in myocardium, we observed no histopathologic effects on the cardiac muscle tissues. In the spleen and regional lymph node tissues, we observed mild lymphopenia in two cases, which probably does not correlate with the vaccine administration. During the examination of liver specimens, some showed mild perivascular and/or sinusoidal hyperemia, which can occur during anaesthesia; however, in pig #3 we detected mild to moderate necrotizing hepatitis. As it was a single case, the relation of cause and effect between vaccination and the hepatic pathology is highly questionable; it is more likely that this pig had a viral/bacterial infection – further examination would be needed to rule this out. We observed a few tubular cysts in the kidney sections, with either flattened or normal cuboidal epithelial cells, or mild hyperemia. Cyst formation in the swine renal cortex is often an accidental finding. In conclusion, we can say that no major pathologies were detected in these specimens, except pig #3.

**SFigure 1.**
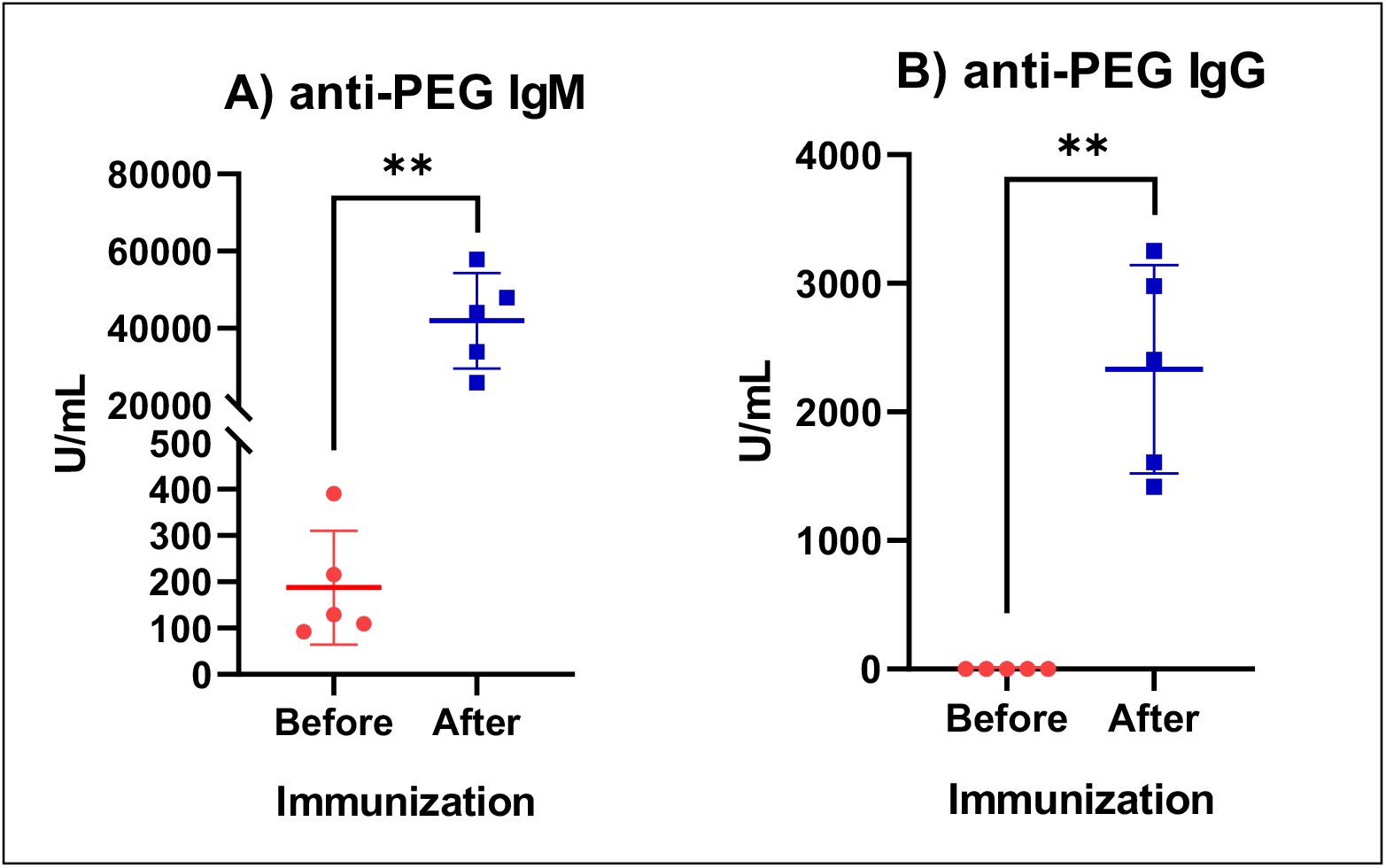
The induction of anti-PEG IgM and IgG by PEGylated doxorubicin-free liposomes. A) anti-PEG IgM and B) anti-PEG IgG one week after i.v. infusion of pigs with Doxebo. Values are mean

